# Stochastic Models for Population Dynamics

**DOI:** 10.1101/031237

**Authors:** Sean X. Sun

## Abstract

Cell growth and division are stochastic processes that exhibit significant amount of cell-to-cell variation and randomness. In order to connect single cell division dynamics with overall cell population, stochastic population models are needed. We summarize the basic concepts, computational approaches and discuss simple applications of this modeling approach to understanding cancer cell population growth as well as population fluctuations in experiments.

## I. Introduction to Stochastic Population Models

One of the simplest experiments is to grow a clonal population of cancer cells, starting with a single cell. Assuming there are no nutrient and space limitations, each cell will undergo a round of symmetric cell division, produces 2 daughter cells and in the process increases the cell population by 1. Careful measurements, however, show that the time between each round of cell division is not a constant, but exhibit significant amount of variation [1]. For example, Fig. 1 shows the collected cell cycle time distribution (also known as *waiting time distribution* for cell division) for human dermal fibroblast (HDF) cells. We see that there is a significant portion of cells that divide at 30hrs, fully 10hrs longer than the average division time of 20hr. Some cells also divide significantly faster. Quantitatively, this means that if we started from a single cell and counted the cell population, *N*, after time *t*, we would obtain a different *N* each time we perform the identical experiment. We can perform the experiment many times and build a statistical distribution of cell population *P*(*N,t*). This distribution contains information about underlying events such as cell division and death. Modeling such a population distribution is the subject of this chapter.

**Fig. 1.**
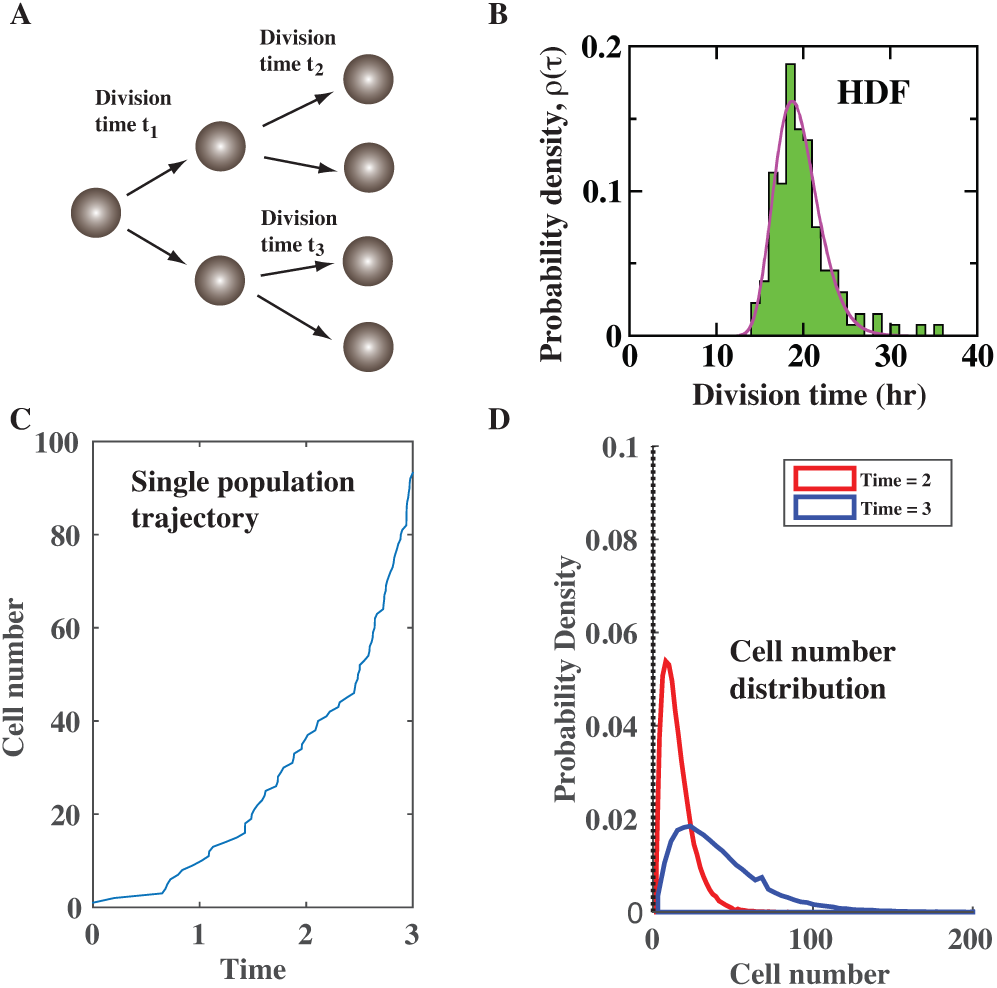
(A) A simple cell growth experiment where each cell undergoes symmetric cell division. The division time is a stochastic variable. (B) Measured distribution of cell division time for HDF cells. The average is close to 20hr, but the variation is large. (C) The number of cells in a single stochastic simulation. The cell number increases stochastically. (D) After simulating many growth trajectories, the cell number distribution shows that there can be a wide variation in the number of cells.

There are many reasons why stochastic population dynamics is important in cancer. First, no tumors are alike, and even with genetically identical cells. The simple fact that cell division is stochastic suggests that tumor size will vary. Second, in a tumor environment, cell division and death depend on many factors, including phenotype of the cell. There is competition as well as cooperation between cells. Therefore the cell population distribution can be complex, and perhaps only rare parts of the distribution will lead to metastasis and cancer progression. These questions cannot be addressed without stochastic population models.

To illustrate how to apply stochastic models to population growth, we revisit the cell population growth problem in the first paragraph. Let us consider how the distribution of cell number can change in time. Here we will assume Markovian dynamics, namely that cells do not have memory about the environment long ago, and only grow according to the current condition. (Note that Markovian dynamics does not mean that there are no population correlations in time. This still can happen because it takes finite time for distributions to evolve.) In this limit, the cell number distribution at *t+Δt* is *P*(*N,t* + Δ*t*) = *u*(*N* − 1)*P*(*N* −1,*t*) + (1 − *uN*)*P*(*N*,*t*), where *u* is the probability of a cell division to occur in Δt. *u(N-1)* is the total probability of *N-1* cells to increase to *N* cells in Δt, since the division event can occur for any of the *N-1* cells. *P(N,t+Δt)* increases when one of *N-1* cell divides and give rise to *N* cells (first term). *(1-uN)* is the probability of no cells divided in time *Δt* (second term). From here, we can obtain the so called Master equation by subtracting both sides by *P(N,t)* and dividing by *Δt*. We obtain a differential equation of the form:

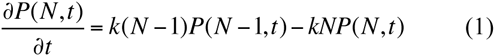
where *k=u/Δt* is the *rate* of cell division. This simple equation allows us to compute the cell number distribution over time. It is worth noting that even for this very simple example, the results show a large variation in the number of cells, as illustrated in Fig. 1. Here, *k=1* and we see that there is substantial probability to observe *N=100* at *t=3,* even though the average number of cells is close to 15.

From the distribution *P(N,t),* we can obtain all the relevant statistical information. For instance, the average cell population is just 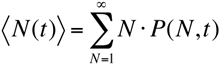. The mean squared variation is 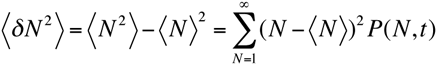. The average population and the variation in population can be measured experimentally, and the data can be used to extract parameters such as the cell division rate, *k*.

Going beyond the simple cell growth example, the same stochastic population framework can be used to examine more interesting scenarios where results maybe surprising. For example, in realistic conditions, there are cell division and cell death. Division and death decisions can also depend on conditions of the cell population such as the population composition or the total population [2]. For example, when space or nutrients becomes limiting, the cell division rate may slow down as the population increases. Thus, *k* would depend on *N*, e.g., *k = k*_0_ − *γN*. In this situation the population can reach a steady state, where there is a cell number distribution that does not change with time. Although microscopically, there continues to be cell division and cell death.

The scenario becomes more interesting when there are several different types of cells that compete, or cooperate in a tumor environment. These interactions between different cell types can introduce complexity and unexpected behavior. In these case, we need to consider more complicated distributions, *P(N,M,t),* where *N* is the number of cell type 1, and *M* is the number of cell type 2. One can then write down Master equations like in Eq. 1 and consider dynamics of the population joint distribution. Here, stochastic population models may predict that multiple compositions of cell populations may exist for the same environment [3]. In an experiment, one may observe abrupt transitions between these different types of population in a random manner. Within the stochastic framework, there are also situations where the population may reach a ‘critical point’ where the population variation becomes very large and 〈*δN*^2^〉diverges [4]. These stochastic population phenomena may have significant implications for cancer, and the mechanisms and consequences are still being examined.

It is also important to contrast the results from stochastic population models with models that consider the average population only. For example, for the simple cell division problem in Fig. 1, the equivalent average population model is *dn/dt=kn,* where *n* = 〈*N*(*t*)〉. The average population model can be derived from the Master equation [5]. One can show that as *n* increases, the population variation becomes small when compared with the average. However, strictly speaking when the average population is small, the evolution of the average population depends on the population variation as well as higher order moments of the full distribution *P(N,t).* Therefore, when the population is small, the average population model in fact fails, and the stochastic population model is a more precise way of describing population change.

## II. Stochastic Simulations for Population Models

For complex populations with several cell types and multiple dependences in cell division and cell death, the full Master equation may be too complex to solve using traditional methods. An alternative approach is to use stochastic simulation methods based on random generation of cell division and cell death events [6]. For the simplest problem discussed in Fig. 1, we can randomly pick a time of cell division according to the measured division time distribution. If there are *N* identical cells, then each cell may undergo replication according to the division time distribution. In this way, we can ‘simulate’ the growing population. To include other events such as cell death, or phenotypic change, multiple events must be randomly sampled according to their waiting time distributions. For exponential waiting time distributions, i.e., 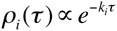 where *i* labels the type of event (death, division, or phenotypic change), we can chose a random number, *r*, and compare with 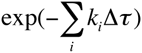 where Δτ is a predefined small time increment. If 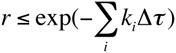, then an event will occur during the time increment. The identity of the event is determined by another random number compared with the relative weight of individual rate parameters 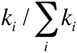. By successively incrementing time and generating random numbers, we can generate stochastic ‘trajectories’ of cell populations over time. By performing the simulation many times, we can collect statistics from the trajectories and obtain the population distributions for the problem at hand. Therefore, the simulation approach is a direct test of the underlying assumptions on the cell division rate, cell death rate and any other rate parameters in the problem.

## III. Quick Guide to the Methods

To implement stochastic population models, one typically assumes a waiting time distribution for all the stochastic events (division, death, phenotypic transformation, etc) in the problem. Commonly, exponential waiting time distributions are assumed, which implies that rates of stochastic events, *k_i_*, are constants in time. Note that *k_i_* can still depend on the current population *N*, or environmental variables. The other typical assumption is that the system is Markovian, i.e., all events occur according to the current population or environment of the system and there is no memory of past system state. It is useful to note that for typical eukaryotic cell division, the waiting time distribution is not exponential and therefore violate the first assumption.

### A. Equations

The equations used in stochastic populations models are different versions of the Master equation in Eq. 1. These equations can be complex, depending on the growth mechanism. For example, if cell division slows down as the population increases, *k = k*_0_ − *γN*, then Eq. 1 becomes:

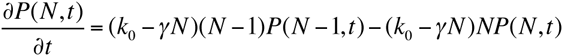

Master equations can be solved using matrix methods. Stochastic simulation should produce the same *P(N,t)* as the Master equation.

### B. Type of settings in which these methods are useful

Stochastic population models are useful for examining underlying mechanisms that generate the cell population composition observe in experiments. By ‘mechanism’, we mean how cells control cell division, cell death, and phenotypic transformation events, and how environmental conditions influence these events. An example of such control is decreasing cell division rate with increasing cell population (when cells entering into a quiescent state). Here, this mechanism implies that *k* = *k*_0_ − *γN*. In complex tumor environments, there are multiple influences on cell division and death, and these influences are critical for determining cell population over time. Stochastic population models are also essential for understanding variations in cell populations seen in population dynamics.

